# Predatory Bacteria can Reduce *Pseudomonas aeruginosa* Induced Corneal Perforation and Proliferation in a Rabbit Keratitis Model

**DOI:** 10.1101/2023.03.15.532777

**Authors:** Eric G. Romanowski, Nicholas A. Stella, Bryn L. Brazile, Kira L. Lathrop, Jonathan M. Franks, Ian A. Sigal, Tami Kim, Mennat Elsayed, Daniel E. Kadouri, Robert M.Q. Shanks

## Abstract

**Purpose:** *Pseudomonas aeruginosa* keratitis is a severe ocular infection that can lead to perforation of the cornea. In this study we evaluated the role of bacterial quorum sensing in generating corneal perforation and bacterial proliferation and tested whether co-injection of the predatory bacteria *Bdellovibrio bacteriovorus* could alter the clinical outcome. *P. aeruginosa* with *lasR* mutations were observed among keratitis isolates from a study collecting samples from India, so an isogenic *lasR* mutant strain of *P. aeruginosa* was included.

**Methods:** Rabbit corneas were intracorneally infected with *P. aeruginosa* strain PA14 or an isogenic Δ*lasR* mutant and co-injected with PBS or *B. bacteriovorus*. After 24 h, eyes were evaluated for clinical signs of infection. Samples were analyzed by scanning electron microscopy, optical coherence tomography, sectioned for histology, and corneas were homogenized for CFU enumeration and for inflammatory cytokines.

**Results:** We observed that 54% of corneas infected by wild-type PA14 presented with a corneal perforation (n=24), whereas only 4% of PA14 infected corneas that were co-infected with *B. bacteriovorus* perforate (n=25). Wild-type *P. aeruginosa* proliferation was reduced 7-fold in the predatory bacteria treated eyes. The Δ*lasR* mutant was less able to proliferate compared to the wild-type, but was largely unaffected by *B. bacteriovorus*.

**Conclusion:** These studies indicate a role for bacterial quorum sensing in the ability of *P. aeruginosa* to proliferate and cause perforation of the rabbit cornea. Additionally, this study suggests that predatory bacteria can reduce the virulence of *P. aeruginosa* in an ocular prophylaxis model.

## 1. Introduction

Predatory bacteria, such as *Bdellovibrio bacteriovorus*, have been suggested as an alternative approach for treatment antibiotic resistant bacterial infections in general [1–6]. *B. bacteriovorus* is a Gram-negative bacterium that is an obligate predator of other Gram-negative bacteria. Given the preponderance of Gram-negative bacteria in causing contact-lens related keratitis, predatory bacteria have been hypothesized as treatments for eye infections [7–9], and are largely non-toxic and non-inflammatory when applied in great numbers to a corneal epithelial cell line and intact and wounded ocular surfaces [7, 10]. These have been tested to prevent bacterial proliferation on the ocular surface with some success and to prevent the development of keratitis in a mouse model using *Escherichia coli* as a pathogen [11, 12].

Unlike *E. coli, Pseudomonas aeruginosa* is the leading cause of contact lens associated keratitis, a blinding infection [13–15]. Antibiotic resistant *P. aeruginosa* keratitis isolates are associated with worse clinical outcomes [16–19]. Previous work showed that *B. bacteriovorus* predation is not influenced by antibiotic resistance status of its prey [3, 5, 7] and is even effective at clearing antibiotic tolerant biofilms [20, 21]. A recent study showed that around 20% of *P. aeruginosa* keratitis isolates isolated in India from 2006-2010 had mutations in the *lasR* gene and that these strains were correlated with worse visual outcomes in patients [22]. The LasR protein is a major regulator of quorum sensing and important in regulation of numerous virulence-associated genes [23].

In this study we evaluated whether *B. bacteriovorus* strain HD100 could act as a “living antibiotic” in a rabbit keratitis infection model using a clade 2 (cytotoxic/ExoU^+^) and highly virulent *P. aeruginosa* strain. This was a proof-of-concept prevention model rather than a treatment model. The study found that the predatory bacteria caused mild-moderate inflammation, but were able to reduce *P. aeruginosa* proliferation and most importantly significantly reduced the frequency of corneal perforation events caused by *P. aeruginosa*. We also observed that an isogenic *P. aeruginosa ΔlasR* mutant was unable to replicate in the rabbit cornea and caused significantly fewer corneal perforations, but was not significantly reduced by *B. bacteriovorus* in the cornea.

## 2. Methods

### 2.1. Bacterial strains and culture

*B. bacteriovorus* HD100 (ATCC 15356) [24] was used for this study. Preparation of *B. bacteriovorus* followed prior reports [10, 12] where *B. bacteriovorus* was incubated with ~10^9^ CFU/ml *E. coli* diaminopimelic acid auxotroph strain WM3064 for 24 h at 30°C. The resulting mixture was passed multiple times through a 0.45-μm Millex®-HV pore-size filter (Millipore, Billerica, MA, USA) to remove remaining prey. Predators were washed with phosphate buffered saline (PBS) and concentrated centrifugation. *B. bacteriovorus* was suspended in PBS to 1.5 × 10^10^ PFU/ml *B. bacteriovorus*.

For a pathogen, the cytotoxic (ExoU+) / clade II strain UCBPP-PA14 (PA14) [25–28] and an isogenic Δ*lasR* mutant were used [29]. These were inoculated from single colonies into LB medium and grown with aeration at 30°C or 37°C. For inoculation, *P. aeruginosa* were diluted in phosphate buffered saline (PBS) to achieve the inoculum (~2-4×10^3^ CFU in 10 μl).

### 2.2. Bacterial keratitis studies

The experiments in this study conformed to the ARVO Statement on the Use of Animals in Ophthalmic and Vision Research and were approved by the University of Pittsburgh’s Institutional Animal Care and Use Committee (IACUC Protocols 15025331 and 18022194).

Female New Zealand White rabbits (1.1-1.4 kg) were obtained from the Oakwood Research Facility through Charles River Laboratories. The rabbits were anesthetized with 40 mg/kg of ketamine and 4 mg/kg of xylazine administered intramuscularly. The corneas of the right eyes only were anesthetized with topical 0.5% proparacaine and injected intrastromally with the 10 μl of *P. aeruginosa* (~2000 CFU) in PBS or PBS alone as a control. Actual inocula for each trial was determined using the EddyJet 2 spiral plating system (Neutec Group Inc., Farmingdale, NY) on 5% trypticase soy agar with 5% sheep’s blood plates. Plates were incubated for ~18-20 h at 37°C and the colonies were enumerated (Flash and Grow colony counting system, Neutec Group, Inc). This was followed by a 25 μl injection of *B. bacteriovorus* in PBS (~4×10^8^) or PBS.

At 24 h post-injection, the eyes were evaluated for ocular signs of inflammation using a slit-lamp according to a modified McDonald-Shadduck grading system [30]. Rabbits were systemically anesthetized with ketamine and xylazine as described above and euthanized with Euthasol solution following the 2020 AVMA Euthanasia Guidelines.

Corneal buttons were harvested using a 10 mm trephine and placed into Lysing Matrix A tubes (MP Biomedicals) containing 1 ml of PBS. The corneas were then homogenized with an MP Fast Prep-24 homogenizer (MP Biomedicals), and the numbers of corneal bacteria were enumerated as described above. The centrifuged and filtered homogenate (Millipore 0.22μm PVDF filter) was frozen for ELISA analysis (IL-1β kit (Sigma-Aldrich RAB1108). MMP9 ELISA) following the manufacturer’s guidelines. Additional corneas were fixed with paraformaldehyde (4%) in PBS, embedded in paraffin, and 5 μm sections were stained with hematoxylin and eosin. Sections were viewed using an Olympus Provis AX-70 microscope and images captured with MagnaFire 2.1 software.

In another set of animals, the whole eyes were removed following euthanasia and stored individually in 25 mm of PBS and kept on ice. Within one hour, their corneas were scanned by optical coherence tomography (OCT) using a Bioptigen Envisu R2210 system with InVivoVue software (Leica Microsystems) following the same general approach described elsewhere [31]. Briefly, the system was modified with a broadband superluminescent diode (Superlum, Dublin, Ireland λ = 870 nm, Δλ = 200 nm, 20,000 A-scans/second). OCT volume scans of the central cornea were acquired using a 10mm telecentric lens for anterior segment imaging and a scan pattern: 6mm x 6mm x 2mm (1000 x 100 x 1024 pixel sampling), with 1 Frame per B-scans, i.e. no repetitions. At least three scans were obtained of each sample and the best quality scan chosen for analysis. The most common issue was a high signal at the cornea apex that reduced visibility directly underneath. Scans of a given eye were completed in less than six minutes. The B-scans were loaded into Fiji for visualization as image volumes and to export stills and movies [32].

### 2.3. Scanning Electron Microscopy

Electron microscopy was performed as previously described [12]. Rabbit corneal buttons were fixed with glutaraldehyde (3%) at room temperature for 24h and washed with PBS. Samples were pinned to prevent curling and post-fixed using osmium tetroxide (1%), dehydrated with ethanol (30–100%), immersed in hexamethyldisilazane and air-dried. Samples were mounted to aluminum stubs and sputter coated with gold/palladium (6 nm). Scanning electron microscopy was performed using a JEOL JSM-6335F scanning electron microscope (3 kV).

### 2.4. In vitro predation assay

*P. aeruginosa* were grown at 37°C in LB broth on a tissue culture rotor, centrifuged and suspended in HEPES buffer (25 mM HEPES with 3 mM MgCl_2_ and 2 mM CaCl_2_). Aliquots of *P. aeruginosa* (0.5 ml, ~1×10^9^ CFU) and *B. bacteriovorus* (0.5 ml, ~2×10^8^ PFU) and 4 ml of HEPES buffer were added to 15 ml conical polypropylene tubes and incubated at 30°C. At 24 h bacteria were serial diluted and plated on LB agar plates to determine surviving *P. aeruginosa*. Log reductions were calculated as compared to a predator free control survival at the same time point.

### 2.5. Statistical analysis

Kruskal-Wallis with Dunn’s post-test was used for non-parametric analysis and ANOVA with Tukey’s post-test or Student’s T-tests for parametric analysis. Contingency analysis was performed with Chi-Square test followed by Fisher’s Exact test for pair-wise comparisons. All analyses were done with GraphPad Prism software.

## 3. Results

### 3.1. Evaluation of the antimicrobial efficacy of *B. bacteriovorus* in a *P. aeruginosa* keratitis model

The ability of the predatory bacteria *B. bacteriovorus* strain HD100 to prevent *P. aeruginosa* proliferation in a bacterial keratitis model was evaluated. *P. aeruginosa* was used because it is a major cause of corneal infection. In this study either 10 μl of *P. aeruginosa* (~2000 CFU) or the PBS vehicle was injected into the corneal stroma (Figure 1A), followed by a second injection of 25 μl of either *B. bacteriovorus* strain HD100 (4×10^8^ PFU) or the PBS vehicle into the same needle track.

**Figure 1.**
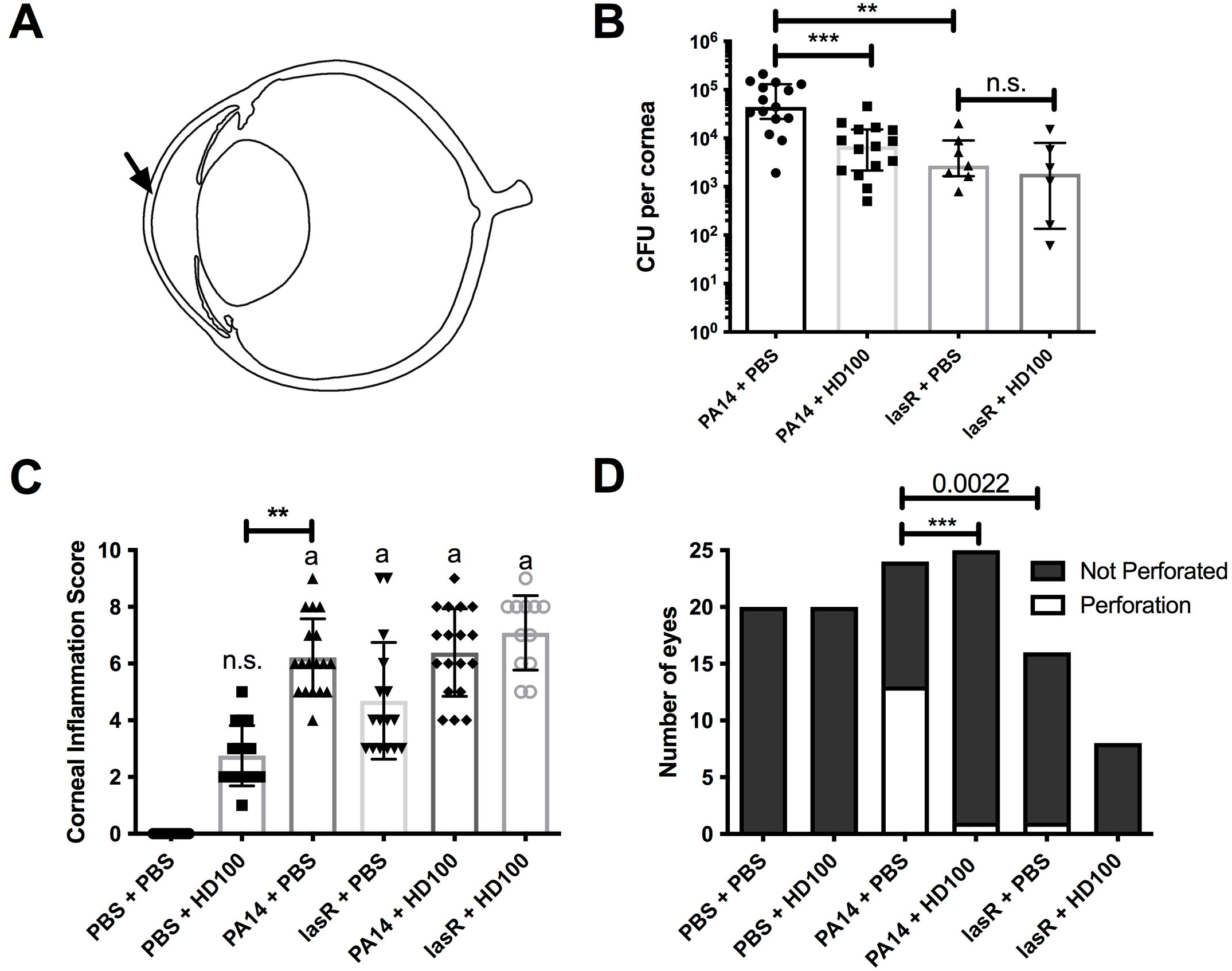
Predatory bacteria and *lasR* mutation reduce *P. aeruginosa* pathogenesis in a rabbit keratitis model. **A.** Diagram of model in which the rabbit corneal stroma (arrow) is injected with *P aeruginosa* strain PA14, isogenicΔ*lasR* mutant, or PBS followed by PBS or *B. bacteriovorus* strain HD100. **B-D**. Each point represents a rabbit. **a** indicates significant difference from control group, p<0.05. n.s. indicates not significantly different than the control group. **, p<0.01; ***, p<0.001.**B**. CFU from corneas at 24 h post-injection. Means and SDs are shown. **C.** Corneal inflammation score based on MacDonnald-Shadduck scoring system. Medians and interquartile ranges are shown and Kruskal-Wallis analysis with Dunn’s post-test was performed. **D.** Corneal perforation frequency. Indicated groups were compared by Fisher’s Exact test.

The virulent *P. aeruginosa* strain UCBPP-PA14 (PA14) [25–28] was injected into the cornea at an average of 1,777±353 CFU/cornea. This strain is a clade 2 strain, also known as a cytotoxic strain, that is ExoU positive and ExoS negative [28]. This strain has not been previously reported in a rabbit keratitis model to our knowledge. PA14 was able to replicate 40-fold to 7.25×10^4^ CFU/cornea by 24 hours (Figure 1B). The eyes were evaluated with a slit lamp and fluorescein staining using a MacDonald-Shadduck-based scoring system [30] and there was a clear increase in corneal inflammation score by 24 h (Figure 1C-D, 2). This led to corneal perforation in just over half of the infected corneas (54%) (Figure 1E, 2).

**Figure 2.**
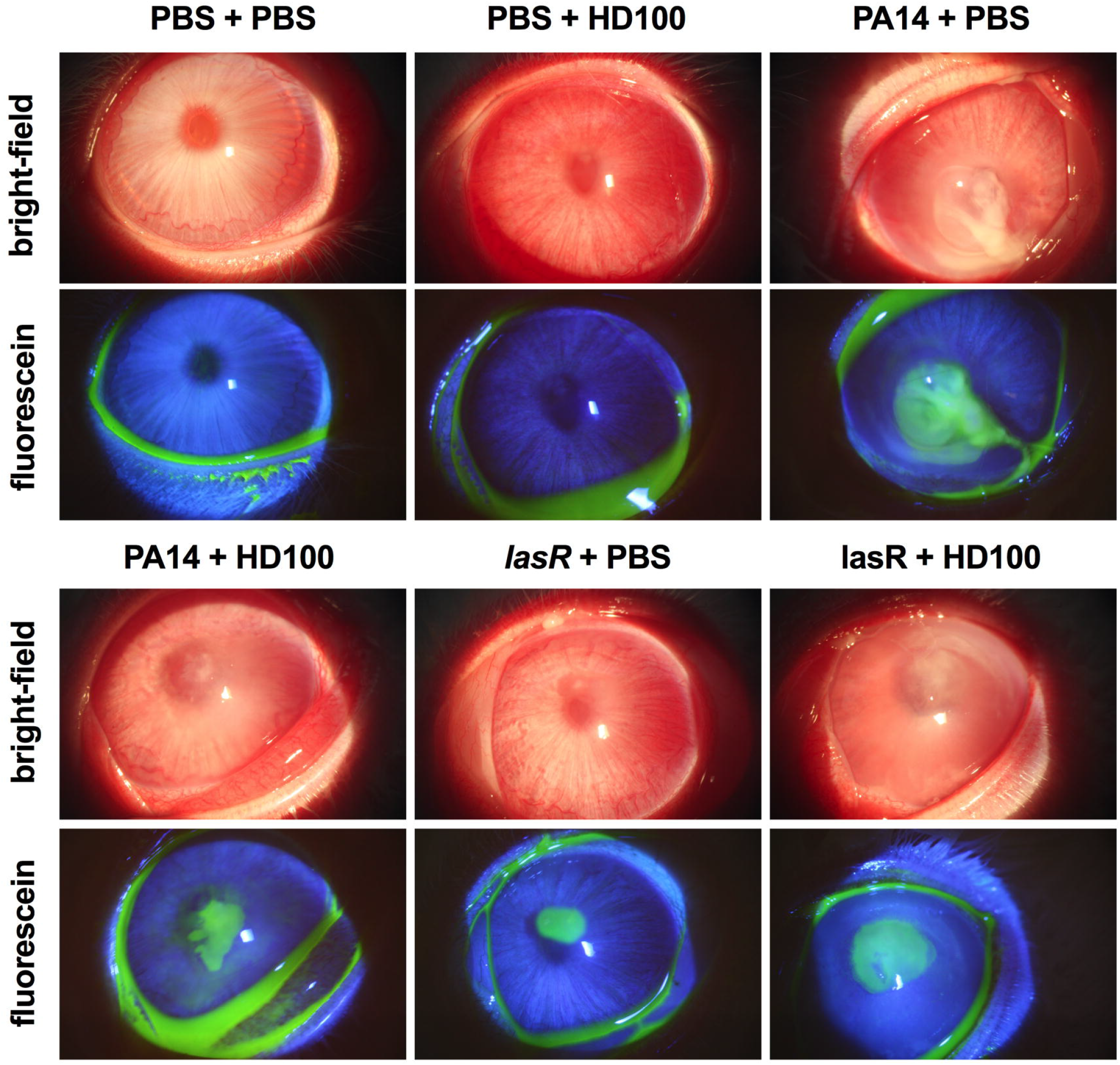
Bright-field microscopy and fluorescein staining of infected eyes. Representative images are shown. Fluorescein staining indicates corneal ulceration and loss of corneal epithelium.

By contrast, *B. bacteriovorus* caused intermediate inflammation compared to PBS and PA14 groups (Figure 1C-E, 2). Moreover, *B. bacteriovorus* injection did not cause corneal perforations (Figure 1D, 2).

*B. bacteriovorus* injected following *P. aeruginosa* correlated with a significant reduction in *P. aeruginosa* proliferation (1.03×10^4^ CFU at 24h, p<0.01) but was still higher than the inoculum. This reduction did not significantly influence the overall corneal score, but was associated with a remarkable reduction in the frequency of corneal perforations (4%, p<0.001) (Figure 1D).

Fluorescein staining and slit lamp examination 24 h post-inoculation revealed minor pathology of the corneas infected with *B. bacteriovorus* that consisted of redness and a small infiltrate that did not cause a fluorescein stained ulcer (Fig 2). By contrast, the *P. aeruginosa* injected eyes showed major purulent discharge from the large central corneal ulcer and anterior chamber effects (Figure 2). Eyes injected with both had an intermediate phenotype similar to those injected with both the Δ*lasR P. aeruginosa* and predatory bacteria. The eyes injected with just the Δ*lasR* mutant bacteria had minimal inflammation, though had a corneal ulcer of moderate size (Figure 2).

Optical coherence tomography (OCT) was done on a subset of eyes following sacrifice of the rabbits at 24h post-inject (Figure 3). This reveals the shape of the cornea and unlike the PBS only injected eye, the *B. bacteriovorus* (HD100) injected eyes had clear inflammation including an infiltrate and overall edema. The loss of the corneal integrity was demonstrated in the *P. aeruginosa* PA14 wild-type injected eyes where the massive infiltrate prevented unobstructed corneal imaging and resulted in shadows. By contrast, the *P. aeruginosa* wild-type and HD100 coinjected eyes had clear damage, ulceration, and infiltrate, but maintained overall shape. The Δ*lasR* mutant with or without HD100 had intermediate phenotypes. Movies depicting OCT volumes of corneal layers are provided in the supplemental information and clearly demonstrate the perforation of the *P. aeruginosa* strain PA14 infected eye (Supplemental movies S1-3).

**Figure 3.**
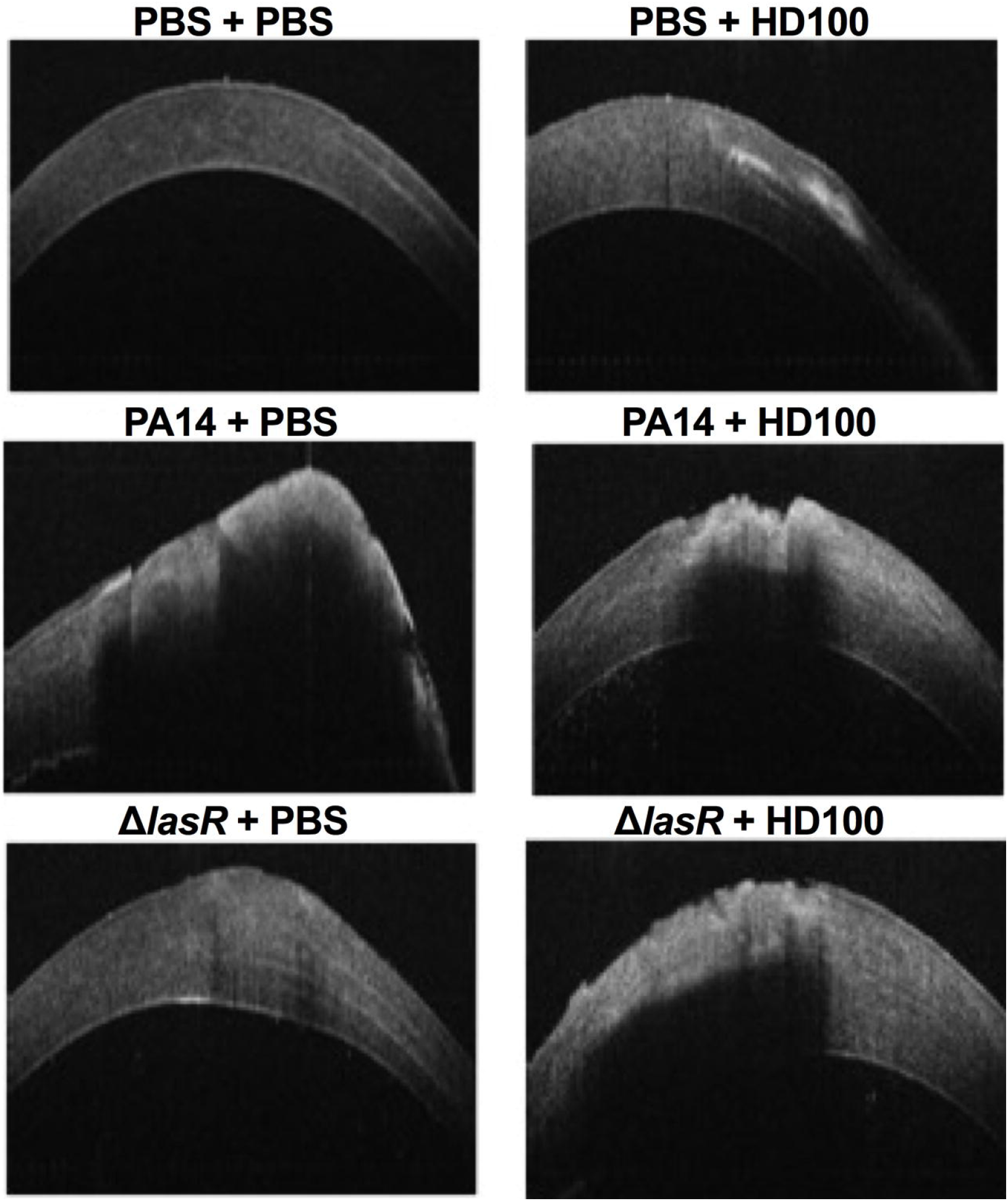
Optical coherence tomography of infected eyes. Representative images are shown. Shadows are due to infiltrates that prevent imaging.

To evaluate whether infiltrates were made by neutrophils, H&E staining of corneas was performed. The histology provided consistent results with the fluorescein staining with respect to maintenance or loss of corneal epithelium (Figure 4). Neutrophils and severe damage of the anterior stroma were observed in corneas infected by PA14 and to a lesser extent by *B. bacteriovorus* (Figure 4 and Figure S1).

**Figure 4.**
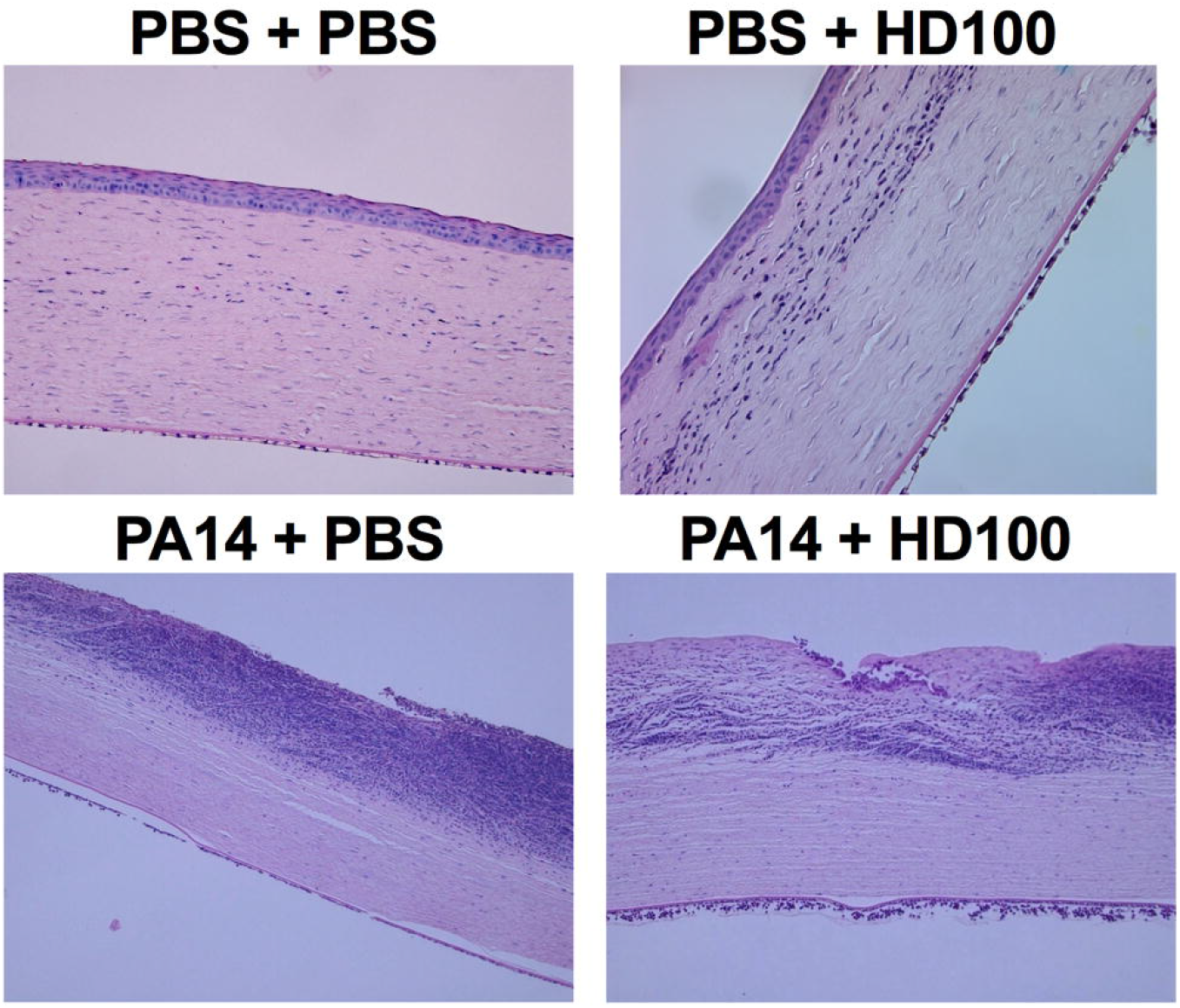
Histological analysis demonstrates neutrophil infiltrates. Representative images are shown. Magnification was 10X. Polymorphonuclear neutrophils are evident from inside the stroma and in at the anterior chamber interface.

Low magnification scanning electron microscopy showed the perforation in the PA14 infected cornea that was absent in the other corneas (Figure 5). Similar to fluorescein results (Figure 2), a clear ulcer was evident in corneas infected with PA14 (PA14 + PBS) and *B. bacteriovorus* (PBS + HD100) (Figure 5). At higher power, red and white blood cells were observed on the ocular surface of most eyes, especially those infected with PA14 (Figure 5).

**Figure 5.**
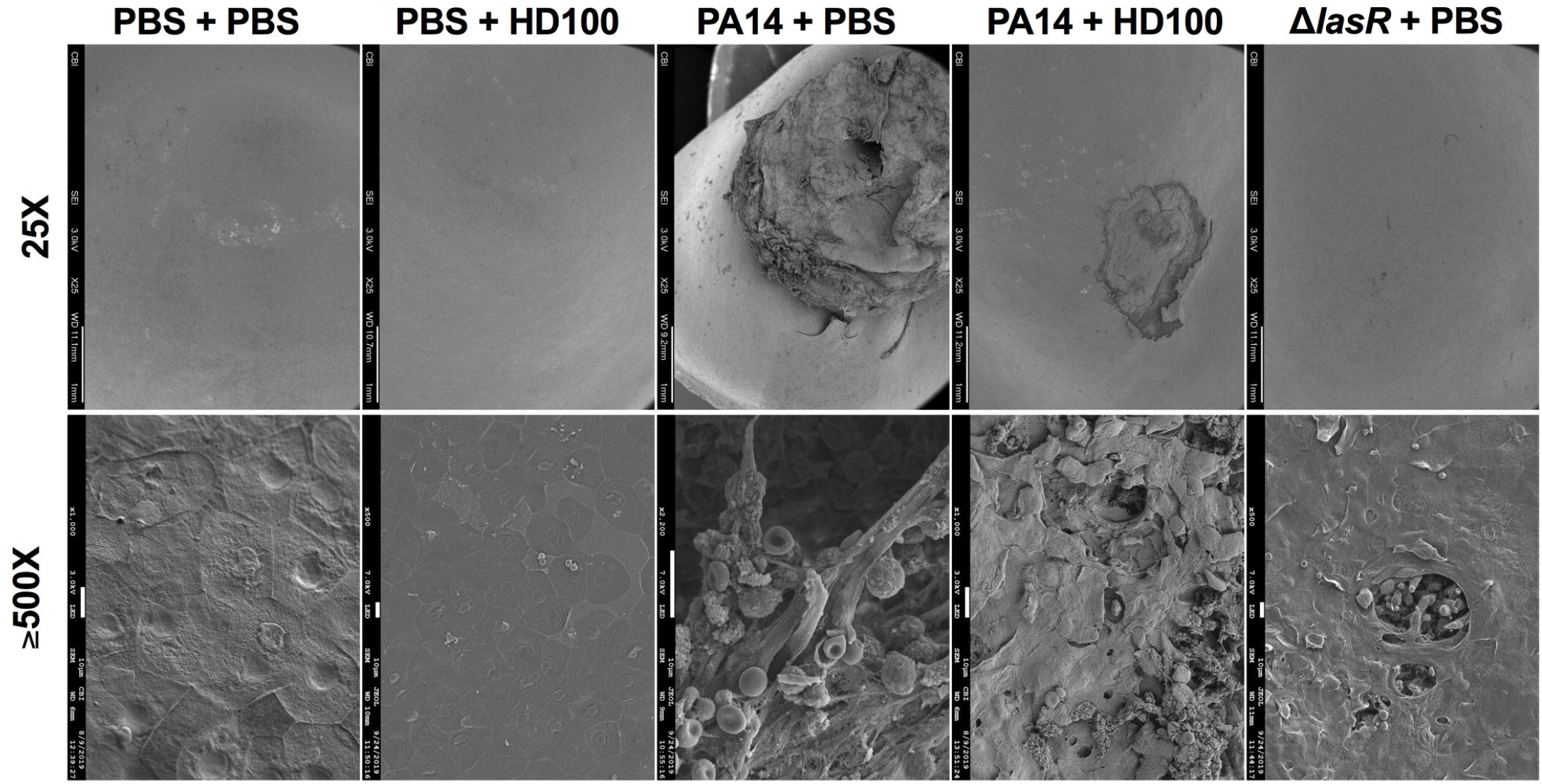
Scanning micrographs of rabbit corneal surfaces. Representative images are shown. Magnification was 25x (top) and ε500x (bottom). The corneal perforation is clearly shown in the PA14 + PBS injected cornea, and a corneal ulcer was evident in the PA14 with HD100 injected cornea.

Consistent with the corneal inflammation MMP9 (92-kDa neutrophil gelatinase or gelatinase B), an enzyme that digests gelatin and collagen types IV and V [33], and the pro-inflammatory cytokine interleukin-1β (IL-1β) were highly induced in corneas infected with PA14, and to a lesser extend by *B. bacteriovorus* (Figure 6). The co-infection of PA14 and *B. bacteriovorus* did not significantly alter these pro-inflammatory markers. However, *B. bacteriovorus* by itself was less inflammatory than PA14 for both MMP9 and IL-1β (p<0.05).

**Figure 6.**
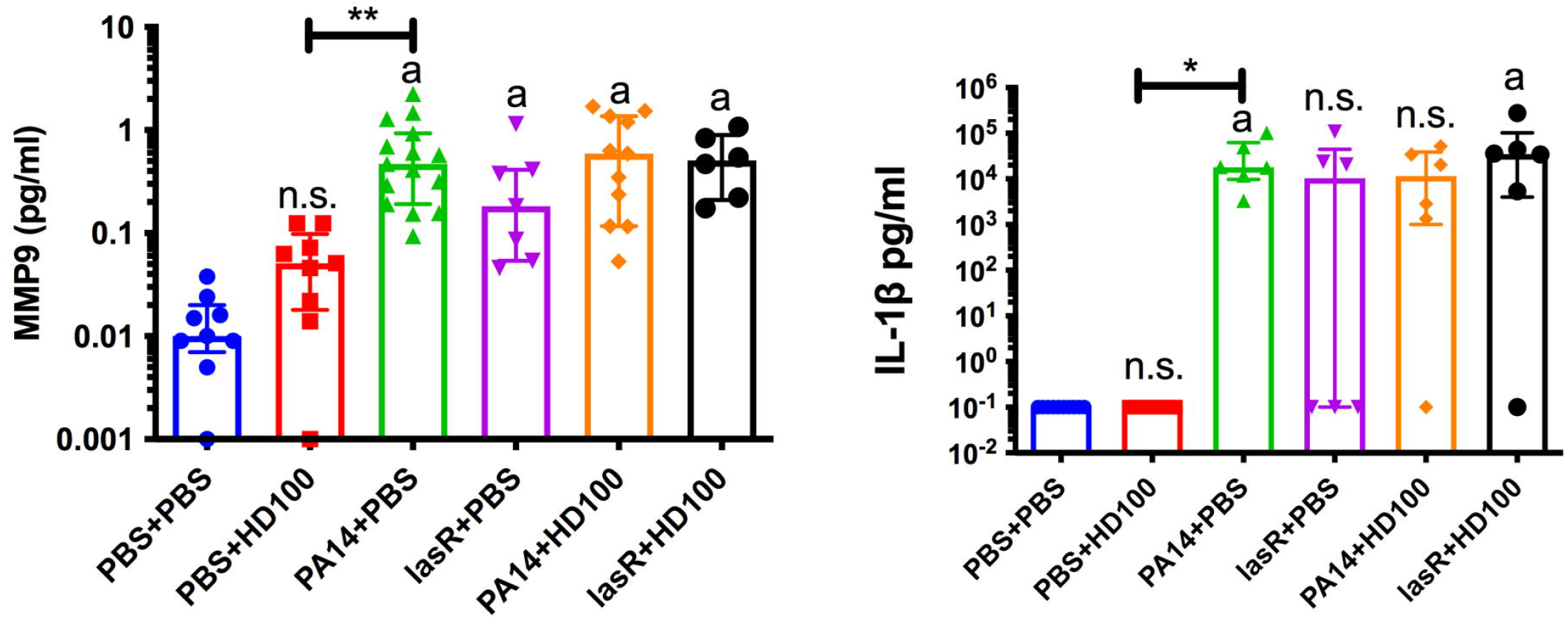
*P. aeruginosa* strain PA14 stimulation of gelatinase B (MMP9) and pro-inflammatory cytokine IL-1® from rabbit corneas. Medians and interquartile ranges of pro-inflammatory cytokines are shown. Each point represents one rabbit. Kruskal-Wallis with Dunn’s post-test was performed, **a** indicates significant difference from control group, p<0.05. n.s. indicates not significantly different than the control group. *, p<0.05; **, p<0.01.

### 3.2. LasR promotes corneal pathogenesis by a cytotoxic *P. aeruginosa* strain

A recent study showed natural *lasR* mutants at high frequency among bacterial keratitis isolates [22]. Therefore, the above experiments were also carried out with an isogenic variant of PA14 with a deletion of the *lasR* quorum sensing regulator. This strain was previously validated by restoring the wild-type allele and is defective in expression of a number of potential virulence associated phenotypes such as protease production [22].

Though slightly more Δ*lasR* mutant bacteria were inoculated compared to the WT (3,805 ± 1703 CFU/cornea, p=0.03), the CFU per cornea at 24 h was significantly lower at 5.9×10^3^ CFU/cornea which is 12-fold lower than wild-type PA14, p<0.01 (Figure 1B). Corneal inflammation scores were also lower, but did not reach significance (p>0.05), (Figure 1C). Nevertheless, the Δ*lasR* mutant was defective in causing corneal perforation compared to the WT (Figure 1D, p=0.002). Histology and SEM analysis demonstrate reduced corneal damage (Figure 2–4), yet the MMP9 and IL-1β levels were similar to the WT (Figure 6). Both the wild-type PA14 and Δ*lasR* mutant was susceptible to *B. bacteriovorus in vitro*. The mean Log_10_ reduction *in vitro* was 4.9±0.05 CFU/ml for PA14 and 4.7±0.15 CFU/ml for Δ*lasR*, p>0.05, n=3. However, the predatory bacteria did not significantly lower the corneal burden. While corneal scores were high, the frequency of perforation was low (0%). However, MMP9 and IL-1β were levels in Δ*lasR* and *B. bacteriovorus* inoculate corneas was indistinguishable from PA14 wild-type infected eyes.

## 4. Discussion

This study demonstrated the severe pathogenesis caused by strain PA14 in a rabbit corneal infection model. 54% of corneas perforated by 24 h limiting the length of the study for ethical reasons. While *B. bacteriovorus* on their own caused some inflammation, they prevented *P. aeruginosa* proliferation and corneal perforation.

In this study, MMP9 was strongly induced by *P. aeruginosa*. MMP9 is a metalloproteinase secreted by neutrophils, macrophages, and corneal epithelial cells [33]. Transcription of the MMP9 gene was strongly induced when corneal cells are exposed to bacteria [34, 35]. MMP9 is involved in remodeling of the corneal basement membrane and regulates corneal healing, but has also been linked with corneal perforation [36, 37]. Tetracyclines inhibit MMP9 activity, and that is one reason that tetracyclines have been given systemically to patients with corneal ulcers or perforations [38]. In animal models tetracyclines promoted wound healing and prevented corneal perforation [39–41]. Our results suggest that MMP9 may contribute to, but is not fully responsible for corneal perforation caused by *P. aeruginosa* as MMP9 levels remained in infected corneas that did not perforate. Other factors such as neutrophil elastase, MMP8, or bacterial proteases such as elastase B, which is positively regulated by LasR may be key contributors to corneal perforation in our model [42, 43].

IL-1 is a major proinflammatory cytokine during bacterial keratitis [44] and promotes neutrophil mediated corneal damage in mouse models of *P. aeruginosa* keratitis [45, 46]. Although a role for IL-1 in preventing corneal perforation in young Swiss mice has also been described [45]. An *in vitro* study showed that keratocytes produced collagenolytic compounds in an IL-1 dependent manner and that collagen degradation could be prevented with an IL-1 inhibitor [47]. IL-1 release from damaged corneal epithelial cells acts as an alarmone and induces increased expression of IL-6 and −8 and reduced barrier function in corneal epithelial cells [48]. Similarly, anti-IL-1β antibody pretreatment reduced MMP9 induction in a mouse (C57Black/6) *P. aeruginosa* keratitis model leading to the overall model that *P. aeruginosa* induced IL-1 induces the expression of MMPs resulting in the destruction of corneal tissue [44].

In our study, predatory bacteria failed to induce IL-1 β production, while this is an unusual for a Gram-negative bacteria in the corneal stroma, *B. bacteriovorus* have unusual physiology that may reduce its ability to activate the immune system. Namely, it has a modified lipopolysaccharide structure that does not activate TLR4 and a membrane sheathed flagellum, so is unlikely to activate TLR5-mediated inflammation

[49, 50]. By contrast, all other infection groups strongly induced IL-1 β that correlated with high levels of MMP9. The intermediate levels of MMP9 induced by *B. bacteriovorus* suggest that there are IL-1-independent mechanisms to induce MMP9. However, only one time point was evaluated, so IL-1 may have been induced at early time point by *B. bacteriovorus*. That said, IL-1 induction by bacterial keratitis was reported to be maintained over the course of several days in a mouse model weakening the likelihood that IL-1 levels were higher at earlier time points [45]. Although the levels of MMP9 measured in the predator group was significantly lower than that measured in the *P. aeruginosa* infected arm, one should consider that the concentration of the predator injected was 5 logs higher (~4×10^8^ vs 2×10^3^ respectively). The low inflammatory stimulating effect of *B. bacteriovorus* seen in this study is in agreement with both *in vitro* [7, 51, 52] and *in vivo* studies that show that pro inflammatory cytokines, including IL-1, are not significantly elevated after exposure to the predator [10, 53–56].

The role of the LasR quorum sensing marker in *P. aeruginosa* keratitis has previously been characterized in mouse models using a clade I/invasive/ExoS+ strain PA01 and similar, but not isogenic *lasR* mutant [57]. Prior work has evaluated the importance of LasR in *P. aeruginosa* keratitis. These studies both used PAO1, an ExoS+ / invasive / clade I strain and a randomly mutated variant of PAO1 that was selected for streptomycin resistance, PAO1s with the *lasR* gene replaced by a tetracycline resistance cassette. The first study demonstrated that in a murine scratch model, 33-fewer *lasR* mutant bacteria were necessary to establish infections in C3H/HeN mice [58]. The second study used the same bacterial strains with a murine scratch keratitis model using BALB/c mice and did not find a difference in the ability of the two bacteria to cause keratitis, however the *lasR* mutant was not as successful in proliferation with ~10-fold lower CFU/cornea being measured [59]. Our study, by contrast, used a ExoU+ / cytotoxic / clade II strain in a rabbit intrastromal injection model. While both the scratch and intrastromal injection models have important limitations, the intrastromal injection model ensures highly similar inoculation numbers, which are important for a treatment study, especially when the *lasR* mutant and wild-type express adhesins at different levels [22]. The results of this study suggest that once past the epithelial barrier the *lasR* mutant is less able to proliferate and this may account for the highly reduced frequency of perforations, additionally, the lower expression of collagen digesting proteases such as elastase B by the *lasR* mutant may also contribute to the highly reduced frequency of corneal perforations [57, 59]. More work is needed to understand the complex role of LasR regulated quorum sensing in *P. aeruginosa* keratitis, nevertheless, this study supports the model that in at least one cytotoxic strain, LasR is required for full levels of pathogenesis.

This proof-of-concept study supports that predatory bacteria can prevent bacterial proliferation and prevent corneal perforation in a keratitis model, however, further study to test efficacy against established infections by multiple types of *P. aeruginosa* clinical isolates are necessary to move the concept of using predatory bacteria as a therapeutic forward.

## 5. Acknowledgements

The authors thank Deborah Hogan at Dartmouth Medical School for strains, Kimberly Brothers and Katherine Davoli for help with animals and histology expertise. This study was funded by National Institutes of Health grants R01EY027331 (to R.S.) and CORE Grant P30 EY08098 to the Department of Ophthalmology. The Eye and Ear Foundation of Pittsburgh and from an unrestricted grant from Research to Prevent Blindness, New York, NY provided additional departmental funding. This work was also funded by the U.S. Army Research Office and the Defense Advanced Research Projects Agency (DARPA) and was accomplished under Cooperative Agreement Number W911NF-15-2-0036 to DEK and RMQS. The views and conclusions contained in this document are those of the authors and should not be interpreted as representing the official policies, either expressed or implied, of the Army Research Office, DARPA, or the U.S. Government. The U.S. Government is authorized to reproduce and distribute reprints for Government purposes notwithstanding any copyright notation hereon.

## 6. Disclosure/Conflict of Interest Statement

The authors have no commercial interests regarding this study.

## Notes

### Competing Interest Statement

The authors have declared no competing interest.

