## Supplementary figures and images for "Predatory Bacteria can Reduce *Pseudomonas aeruginosa* Induced Corneal Perforation and Proliferation in a Rabbit Keratitis Model"

### Figure S1 copy.tiff

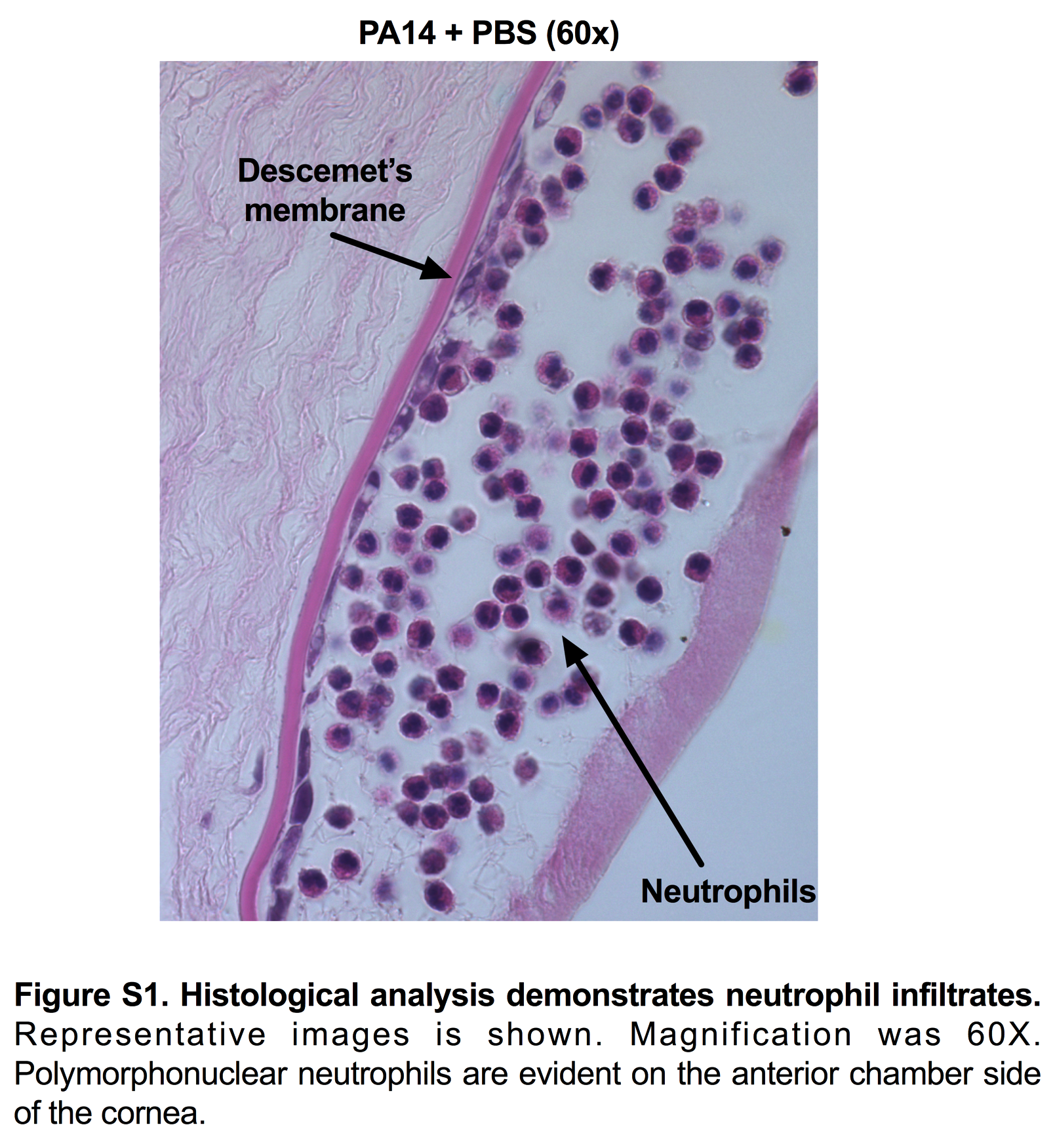
